# Rapid Inactivation of Severe Acute Respiratory Syndrome Coronavirus 2 (SARS-CoV-2) by Tungsten Trioxide-Based (WO_3_) Photocatalysis

**DOI:** 10.1101/2020.08.01.232199

**Authors:** Silvia Ghezzi, Isabel Pagani, Guido Poli, Stefano Perboni, Elisa Vicenzi

**Affiliations:** Viral Pathogenesis and Biosafety Unit, IRCCS San Raffaele Scientific Institute, Milan, Italy; Vita-Salute San Raffaele University School of Medicine, Milan, Italy; Nanohub, Milan, Italy

## Abstract

Severe acute respiratory syndrome coronavirus-2 (SARS-CoV-2), the etiological agent of coronavirus disease 2019 (COVID-19), is transmitted person-to-person via respiratory droplets and, likely, via smaller droplet nuclei light enough to remain suspended in the air for hours and contaminate surfaces particularly in indoor conditions. Thus, effective measures are needed to prevent SARS-CoV-2 transmission in indoor environments. In this regard, we have investigated whether a system based on a filter combining Tungsten Trioxide-Based (WO_3_) photocatalysis and an antiviral fabric treated-copper nanocluster could inactivate SARS-CoV-2. To this purpose, an infectious SARS-CoV-2 suspension was introduced in the upper opening of a closed cylinder containing a WO_3_ filter and a light-based system that activates WO_3_ and the antiviral fabric. From the bottom exit, aliquots of fluid were collected every 10 min (up to 60 min) and tested for their infectivity by means of a viral plaque assay in Vero cells whereas, in parallel, the viral RNA content was measured by quantitative PCR (qPCR). As we have previously shown for SARS-CoV, a 1:1,000 ratio of plaque forming units (PFU) vs. viral RNA copies was observed also for SARS-CoV-2. After 10 min, the infectious viral content was already decreased by 98.2% reaching 100% inactivation after 30 min whereas the SARS-CoV-2 RNA load was decreased of 1.5 log_10_ after 30 min. Thus, in spite of only a partial decrease of viral RNA, SARS-CoV-2 infectivity was completely abolished by the WO_3_ photocatalysis system by 30 min. These results support the hypothesis that this system could be exploited to achieve SARS-CoV-2 inactivation in indoor environments.

## Introduction

At the end of 2019, a novel severe respiratory disease (coronavirus disease 2019, COVID-19) emerged in Wuhan, China and has since become pandemic in a few months, with more than 17 million of people infected worldwide as of today ((https://gisanddata.maps.arcgis.com/apps/opsdashboard/index.html#/bda7594740fd40299423467b48e9ecf6). COVID-19 is caused by a novel coronavirus called severe acute respiratory syndrome (SARS) CoV-2 to distinguish it from SARS-CoV that emerged in Guangdong province in China in 2003 and caused the severe clinical condition known as SARS. Like SARS-CoV, SARS-CoV-2 causes a severe interstitial pneumonia that can lead to acute respiratory distress syndrome (ARDS) and death [1]. However, unlike SARS-CoV, SARS-CoV-2 can also cause a multi-organ disease with hypercoagulation but also mild symptoms limited to the infection of the upper respiratory tract [2]. Indeed, high viral loads have been detected in the nasal swabs even in the presence of mild symptoms or in asymptomatic individuals [3–5] who can shed and transmit the infection while asymptomatic.

SARS-CoV-2 route of transmission is not yet completely defined as airborne transmission is still debatable [6]. Droplets are expelled when a person speaks, particularly with a loud voice [7], coughs and sneezes [8] whereas aerosols originate from dissemination of droplet nuclei [9]. Both droplets and aerosols have been shown to contain SARS-CoV-2 suggesting that they are a potential source of infectious virus although virus infectivity has not been yet determined [10]. As droplets are classically described as larger entities (>5 μm) as compared with aerosols (<5 μm), they quickly drop to the surfaces and ground by force of gravity whereas aerosols remain suspended in the air for a longer time [11]. Indeed, experimental studies have shown that SARS-CoV-2 virions can remain infectious in aerosol for hours and on inert surfaces up to days [12]. Furthermore, during SARS-CoV pandemic in 2003, a major route of transmission was identified in aerosols generated in the sewing systems as observed in an apartment building in Hong Kong, suggesting that not only infected droplets, but also droplet nuclei, can be a source of infectious virus [13]. As most secondary cases have been reported in indoor environments [6], the need for systems that inactivate infectious virions present in droplets and droplets nuclei has become a priority to prevent spreading infection.

Virus inactivation by physical means has been extensively studied, as reviewed in [14]. In particular, photocatalysis is the natural phenomenon by which a photocatalyst accelerates the speed of a chemical reaction through the action of either natural or artificial light [15]. The main applications use titanium dioxide (TiO_2_)-based photocatalysts that need to be exposed to UV light in order to be activated [16]. In this regard, the development of a new Tungsten Trioxide-Based (WO_3_)-based photocatalyst has significantly increased the effectiveness of photocatalysis and eliminated the need of UV light irradiation [17]. When exposed to light in the visible spectrum, WO_3_ absorbs and converts light energy into electrons and electron gaps. WO_3_ reacts with water (air humidity) and oxygen to create hydroxyl (OH-) and superoxide anions (O_2_-) [18]. Billions of these reactive oxygen intermediates (ROI) are generated and can damage membranes of bacteria, cells and tissues [19]. The ultimate result is an effective decomposition of microorganisms like viruses and bacteria, organic and inorganic pollutants, nitrogen oxides, poly-condensed aromatics, sulfur dioxide, carbon monoxide, formaldehyde, methanol, ethanol, benzene, ethylbenzene and other Volatile Organic Compounds (VOC) [20]. The strong oxidative effect of WO_3_ tungsten trioxide photocatalyst provides the rationale to explore it as a disinfectant of air and solid surfaces. Although many studies have been reported on photocatalytic inactivation of bacteria, however only a few studies have addressed virus inactivation [21].

Here, we have evaluated whether a WO_3_ based photocatalyst system could interfere with SARS-CoV-2 infectivity.

## Methods

The photocatalytic system used in this study is in liquid phase and relies on the combination of two elements developed in order to improve SARS-CoV-2 inactivation: a metallic mesh filter coated with WO_3_ and a cotton fabric soaked in a metallic nanocluster based (CuNh) on a copper solution (colloidal suspension).

The SARS-CoV-2 stock (GISAID accession ID: EPI_ISL_413489) was diluted 1:100 to obtain 80 ml of viral suspension with an infectious titer of 1.7×10^4^ plaque forming units (PFU)/ml. The viral suspension was introduced into the device from its top and aliquots were collected at the bottom every 10 min up to 60 min. The collected viral suspension was then tested for the presence of infectious virus as determined by a previously optimized plaque assay on Vero cells [22] and quantification of viral RNA by real-time PCR, described below.

### SARS-CoV-2 plaque assay

Vero cells were seeded at 1.5×10^6^ cell/well in 6-well plates in Eagle’s Minimum Essential Medium (EMEM) supplemented with 10% fetal calf serum (complete medium). Twenty-four h later, 1:10 serial dilutions of virus containing suspension collected at various times after processing by the device were incubated with Vero cells for 60 min. Cell supernatants were then discarded and 1% methylcellulose (1.5 ml/well) dissolved in complete medium was added to each well. After 3 days, cells were fixed with formaldehyde/PBS solution (6%) and stained with crystal violet (1%; Sigma Chemical Corp.) in 70% methanol. Viral plaques were counted under a stereoscopic microscope (SMZ-1500, Nikon), as published [22]. Viral titers were expressed as PFU/ml.

### SARS-CoV-2 quantitative PCR

Viral RNA was extracted from the viral suspension collected at different time points after processing by the device. Viral RNA was extracted by using QIAamp Viral RNA Mini Kit (Qiagen). Real-time quantitative PCR (qPCR) for the nucleocapsid (N) gene was next performed to determine the viral RNA copies present after inactivation. The viral RNA quantification was carried out with the Quanty COVID-19 Kit (Clonit, Milan, Italy) that includes a reference curve of viral RNA at known copy number with a 7500 Fast Real-Time PCR System (Applied Biosystems).

### Statistical analysis

Prism GraphPad software v. 8.0 (www.graphpad.com) was used for all statistical analyses. Comparison among groups were performed using the one-way analysis of variance (ANOVA) and the Bonferroni’s multiple comparison test. Comparison between two homogenous groups was performed by a paired t-test.

## Results and Discussion

The kinetics of infectious virus, as expressed in PFU/ml, indicates that, after 10 min, the WO_3_ device inactivated SARS-CoV-2 infectious titers by 98.2% and reached 100% inactivation after 30 min **(Figure 1**).

**Figure 1.**
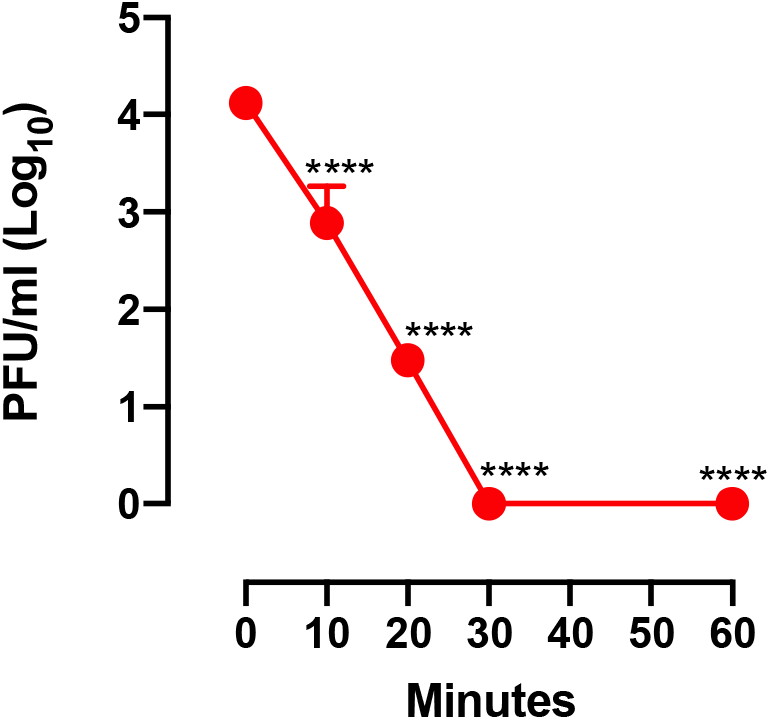
Kinetic of SARS-CoV-2 infectivity inactivation by WO3 treatment. Viral titers were determined by a plaque assay in Vero cells prior to introduction of the SARS-CoV2 into the device and after 10 min of treatment up to 60 minutes. Mean values of three independent experiments are shown. **** indicate a p value < 0.0001 as determined by one-way ANOVA with Bonferroni’s correction.

We next evaluated the amount of viral RNA by qPCR; as shown in **Figure 2A,** the levels of viral RNA in the inoculum were ca. 1,000-fold higher than the infectious titers measured in PFU/ml consistently with previous observations with SARS-CoV [23]. Indeed, the WO_3_ inactivation system progressively and significantly reduced the amount of detectable viral RNA **(Figure 2B),** although not as efficiently as in the case of PFU inactivation.

**Figure 2.**
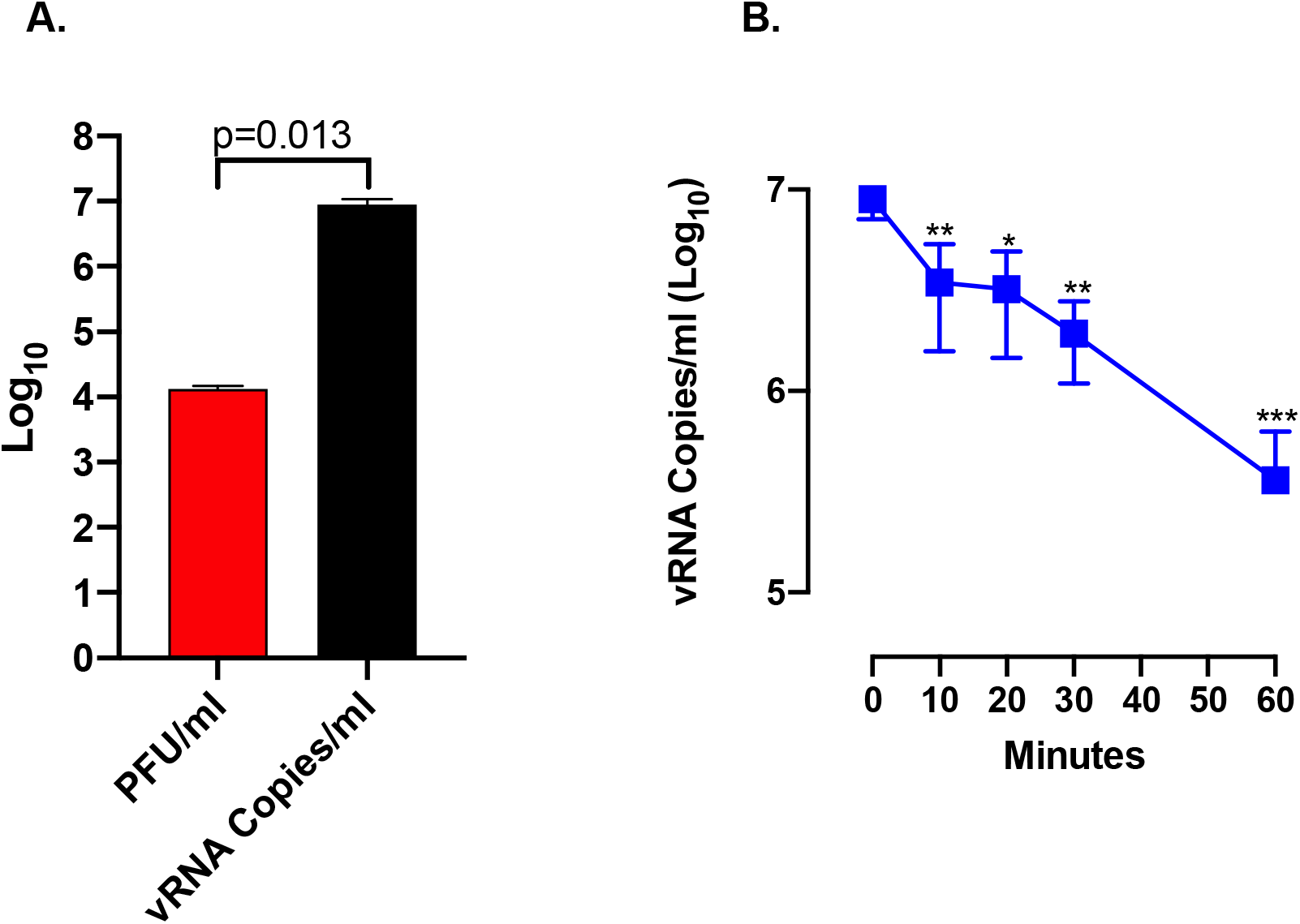
Kinetic of SARS-CoV-2 RNA inactivation by WO3 treatment. **A.** Viral titer of input virus as determined by the plaque assay (red bar) and qPCR (black bar). **B.** Kinetics of viral RNA inactivation by WO3 treatment as determined by qPCR. * indicates a p value < 0.05, ** indicate a p value < 0.01 as determined by one-way ANOVA with Bonferroni’s correction.

Thus, our study supports the hypothesis that a technological device could be potentially exploited for the efficient elimination of infectious SARS-CoV-2, and potentially of other viruses, from the air particularly in close environments where people live, work and spend their time.

The WO_3_ photocatalyst generates a large number of ROI that react very rapidly and efficiently with proteins, lipids and nucleic acids [24]. The destruction of viral proteins, particularly the spike protein that protrudes out of the envelope and the other envelope proteins (i.e. envelope (E) and membrane (M)) likely explains the rapid inactivation of SARS-CoV-2 infectivity. It is tentative to speculate that the generated ROI damage the virion envelope causing the release of genomic RNA, the internal component of viral particles. However, this phenomenon is less efficient than the destruction of the viral protein as genomic RNA is well preserved inside the virion packaged by the tightly bound nucleocapsid protein that likely exerts a shield effect [25].

In comparison to other devices, such as those based on UV light, this system has the advantage to be used safely in the presence of people. All reactions (e.g. virus inactivation and disintegration of other substances) take place in the filter without the release of substances that could be potential hazardous to human and animal health. Furthermore, this system has minimum maintenance requirements and low electricity consumption. Although the test was carried out in liquid solution through SARS-CoV-2 contact with the photocatalytic filter and antiviral tissue, it is highly likely that the same results would be obtained with an air treatment device that uses the same filtration system in indoor environments.

In conclusion, this system has the potential to significantly reduce SARS-CoV-2 infectious titer in the air of indoor environments and, consequently, the contamination of inert surfaces thus contributing to a more general containment of the pandemics in synergy with social distancing and individual measures of protection and hygiene.

